# The distribution of connective tissue in humans – Reconstruction and classification

**DOI:** 10.1101/2025.02.18.638801

**Authors:** Heiko Stark, Julian Sartori

## Abstract

This study aimed to measure human connective tissue distribution and characterise material properties. We used, therefore, the existing data sets of a male and female body donor from the Visible Human Project (VHP). In the first step, collagen-containing structures were segmented and reconstructed using digital image processing. In addition, several properties were characterised and analysed using computational methods. The percentage of fibrous components within the whole body was highest in the thigh. The results of the local orientation tensors showed that individual fascicles and subtendons can be resolved when isolated but not within compact regions of connective tissues.

## Background & Summary

Surprisingly, humans consist of 60 to 65% water and still have a stable form. Connective tissue, which holds individual cells, organs, and body parts together, plays a unique role here^1–3^. It contributes a significant proportion of protein mass in the body (primary component collagen, 25 to 35%)^4^. Most of the water is present in the body in liquid form or with a higher viscosity (e.g. through hyaluronic acid). In order to stabilise the body, it is organised into compartments. The smallest compartments are located at the cellular level and ensure the integrity of individual cells through the extracellular matrix (ECM). In addition, connective tissue forms unique substructures and compartments for organs (muscles, brain, intestines, etc.) and connects them. The skin is the largest structure that ultimately separates us from the environment.

Collagen forms the ECM of many tissues, often together with other structural proteins such as elastin or biominerals such as bone apatite. Due to this composite character of the ECM, it is essential to know its composition and the distribution and orientation of the components^5,6^. The analogy of reinforced concrete emphasises this necessity. Here, a pressure-resistant and tension-resistant material were combined, resulting in a new material with both properties. As with steel in reinforced concrete, the orientation of collagen fibres (called anisotropy) is essential for the biomechanical characterisation of collagen-containing structures^7^. It develops mainly during ontogenesis and is determined by the principal direction of stress^8,9^. This is important because it is challenging to recreate a comparable structure, for example, if it has been injured or was separated in surgery^10^. This must be taken into account in simulations that investigate surgical changes. The stiffness can differ significantly in the fibre direction than in the directions orthogonal to the fibres^11,12^.

In this study, the connective tissue could be reconstructed across all organs. In addition, material properties were characterised and analysed using image processing methods. The data sets obtained in this way can serve as a basis for simulations and as an overview map for studies investigating details of the connective tissue structure. These can include simulations of the musculature and the bone apparatus. For example, it is known that when a muscle contracts, it not only actively shortens but also transmits lateral forces^13,14^. This is shown by bulging the biceps brachii muscle during a contraction. Connective tissue structures largely transmit these lateral forces and must be considered. However, a detailed description is also necessary for structures with a high proportion of connective tissue. For the knee, for example, Dhaher et al. showed that integrating connective tissue into a model can reveal new biomechanical interactions^15^.

### Aim

This study aimed to measure the distribution of human connective tissue and characterise material properties. However, its primary purpose was to create a database to extend existing and future biomechanical models.

## Methods

### Visible Human Project (VHP)

For this study, we used the existing data sets of a male and female body donor (♂:39 years, 90.26 kg, 1.88 m; ♀:59 years, 88 kg, 1.71 m) from the Visible Human Project® (VHP). The data sets were digitised RGB colour images of cryo-sections with a resolution of ♂:0.144×0.144×1 mm^3^ and ♀:0.144×0.144×0.33 mm^3 16–18^. Body donors were shock-frozen to prepare the cryo-sectioning and divided into four craniocaudal blocks. The blocks were then removed layer by layer, and the surface was photographed. As a result of the quartering, some cross-sections were missing or only contained fragments. This was corrected in a later step.

To use the images from the VHP, the cross-sections first had to be correctly aligned with each other. This was done automatically and manually refined using the software ‘Imaris’ (Oxford Instruments, UK, URL: https://imaris.oxinst.com). In addition, the aligned images were cleaned of artefacts using the software ‘Imagexd’ (Heiko Stark, Jena, Germany, URL: https://starkrats.de), and the missing cross-sections between the blocks were filled by interpolation (fig. 1A).

**Figure 1.**
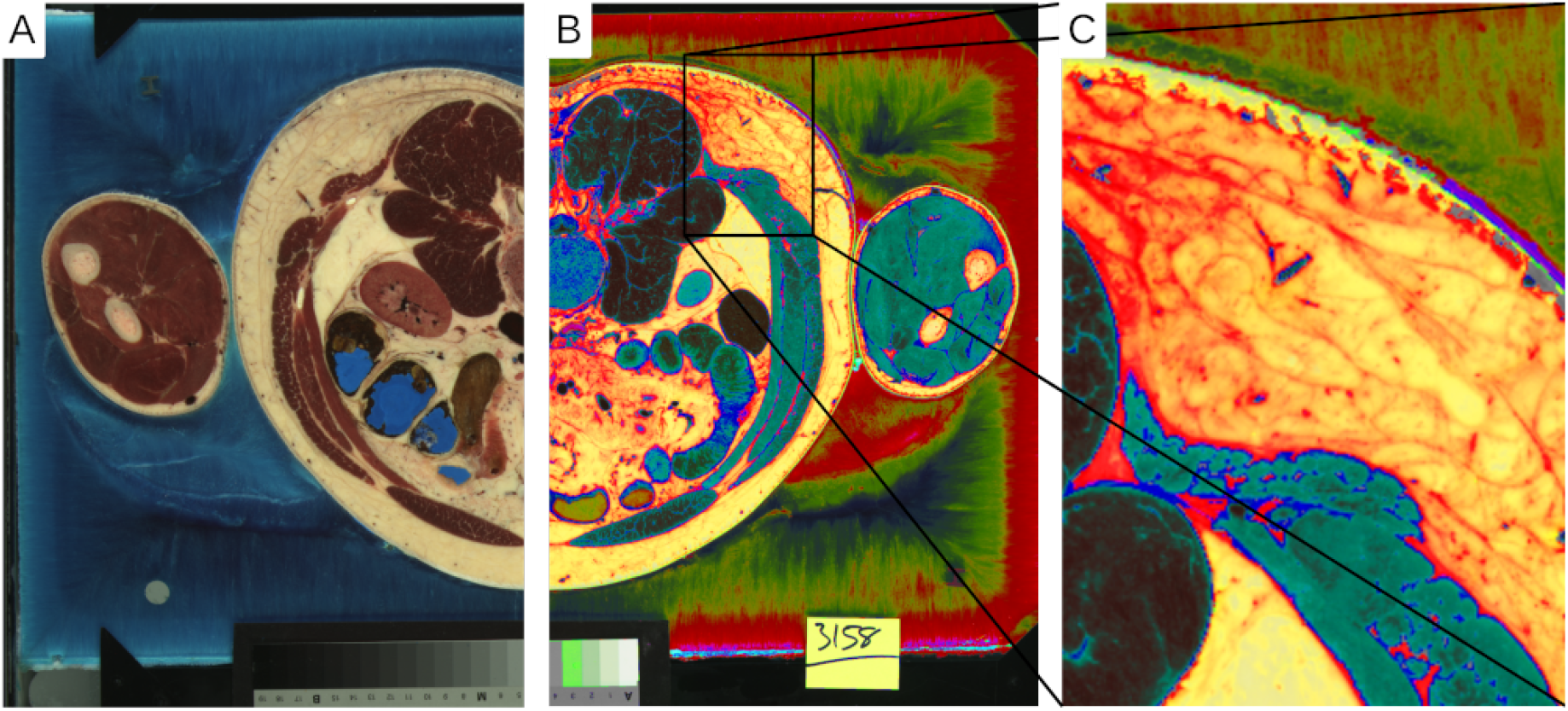
A) Cross-section from the original data set of the Visible Human Project at the level of L3. B) The digitally processed image is based on the RGB information of the connective tissue. C) Closer view from B.

### Image stacks

In the second step, collagen-containing structures were segmented and reconstructed using digital image processing. The approach used the fact that collagen-containing connective tissue had unique colour values in the RGB colour space that could be extracted by a transformation of the colouring (fig. 1B). Furthermore, the colours of connective tissue surrounding the organs and connective tissue embedded in subcutaneous fatty tissue were slightly different (fig. 1C). These values were segmented and further processed to create a topographic model using the software ‘Imagexd’ (fig. 2A-D). Several spectral colours could be considered in the future, which would probably allow an even finer segmentation.

**Figure 2.**
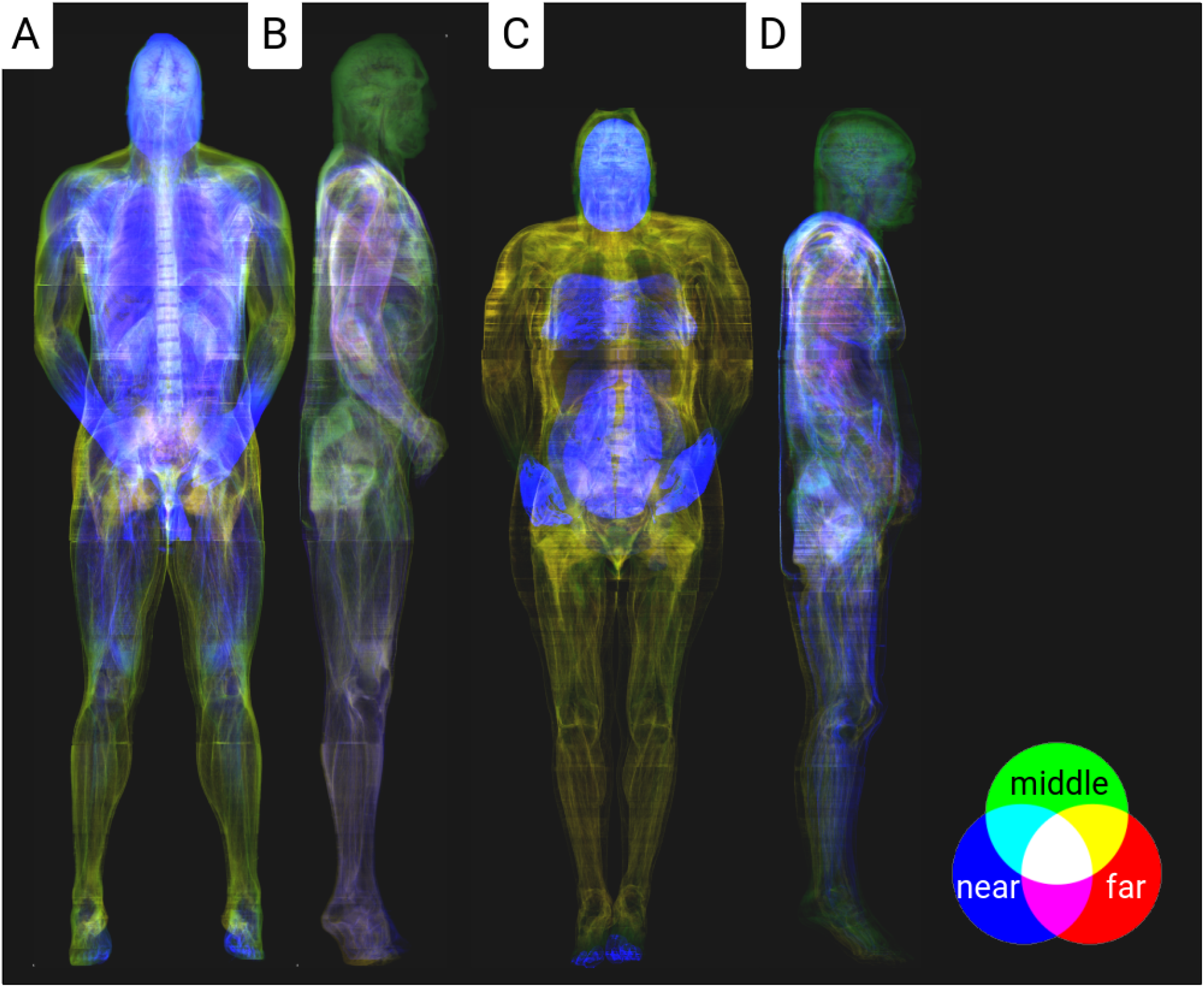
Coloured X-ray visualisation of the connective tissue for the male and female body donor. The colours reflect the depth information of the local connective tissue portions, whereby overlapping colours are averaged.

### Scalar field – density & thickness calculation

The thickness of a material is crucial for many issues. For example, the indenter strength or tissue shell can be estimated this way. Based on the data, the thickness of the connective tissue structures was further calculated using the software ‘Imagexd’. First, the image stacks were converted into isometric voxels (♂:0.5×0.5×0.5 mm; ♀:0.33×0.33×0.33 mm) to simplify the thickness determination. Then, all orientation directions around each voxel with connective tissue content were examined, and the minimum thickness was determined:

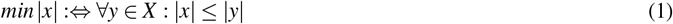

These data were stored in a three-dimensional stack with a 0-32 mm value range and statistically analysed (table 1).

**Table 1.** Statistics of the thickness measurement for selected body regions of the male and female body donors. The mean values of the joint regions in the extremities are highlighted in bold.

| Regio | Male |  | Female |  |
| --- | --- | --- | --- | --- |
|  | Mean | SD | Mean | SD |
| capitis | 5.58 mm | 3.63 mm | 2.64 mm | 2.12 mm |
| cervicales | 5.04 mm | 3.66 mm | 3.09 mm | 2.49 mm |
| thoracales | 3.92 mm | 3.51 mm | 2.48 mm | 2.56 mm |
| abdominales | 2.38 mm | 2.21 mm | 1.83 mm | 1.64 mm |
| perinealis | 3.83 mm | 3.21 mm | 2.02 mm | 1.71 mm |
| deltoidea | **5.90 mm** | 4.18 mm | **2.36 mm** | 1.76 mm |
| brachialis | 3.68 mm | 3.10 mm | 1.89 mm | 1.60 mm |
| cubitalis | **3.52 mm** | 2.60 mm | **2.66 mm** | 3.06 mm |
| antebrachialis | 2.92 mm | 2.41 mm | 1.27 mm | 1.00 mm |
| manus | **3.01 mm** | 2.39 mm | **1.62 mm** | 1.39 mm |
| coxae | **3.71 mm** | 4.03 mm | **1.78 mm** | 1.63 mm |
| femoris | 1.70 mm | 1.81 mm | 1.60 mm | 1.59 mm |
| genus | **2.67 mm** | 2.49 mm | **1.37 mm** | 1.16 mm |
| cruris | 2.18 mm | 1.77 mm | 1.23 mm | 1.02 mm |
| pedis | **2.77 mm** | 2.18 mm | **1.17 mm** | 0.91 mm |

### Tensor field – orientation calculation

To obtain a statistic about the anisotropy of the material, we used a new technique to determine the directions of fascicular structures and the classification of the structures. This was originally developed for a muscular study but generally applies to all fibrous structures^19,20^. A modified method was also applied to the existing isometric voxel data set using ‘Imagexd’. It takes advantage of the fact that local orientation tensors can be calculated from the spatial density distribution. This is a mathematical description of the local anisotropy expressed in a symmetrical 3×3 matrix. The same method was used by Kupczik et al. (2015) and Dickinson et al. (2018) to determine the local vector from the number of possible orientation directions *X*, and it is used to fill the matrix^19,20^. However, in addition, local vectors *x*_*i*_ were summarized using a dyadic product in the respective matrix *A*:

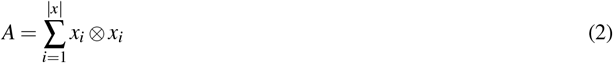

This matrix could be used to determine the three main directional axes (eigenvectors) and their anisotropy strength (eigenvalues). The eigenvalues characterise the tissue components in terms of their structure (table 2). There was only one strongest eigenvector for the fibrous parts and two strong eigenvectors for the planar parts. If all three eigenvectors were equally strong, the structures were classified as a spatially homogeneous collection of connective tissue. This classification could also be colour-coded using the Westin measures *c*_*l*_, *c*_*p*_, and *c*_*s*_ (figs. 3 and 4)^21,22^.

**Table 2.** Statistics on the distribution of the structural proportions for the connective tissue volume of selected body regions of the male and female body donors. The highest structural proportions are highlighted in bold.

| Regio | Male |  |  |  | Female |  |  |  |
| --- | --- | --- | --- | --- | --- | --- | --- | --- |
|  | Volume | Fibrous | Planar | Spherical | Volume | Fibrous | Planar | Spherical |
| capitis | 4.65 dm<sup>3</sup> | 15.49 % | 21.43 % | **50.95 %** | 4.93 dm<sup>3</sup> | 19.81 % | 26.03 % | **31.85 %** |
| cervicales | 0.79 dm<sup>3</sup> | 10.63 % | 18.86 % | **28.77 %** | 0.51 dm<sup>3</sup> | 14.11 % | 19.51 % | **25.94 %** |
| thoracales | 16.50 dm<sup>3</sup> | 18.03 % | 17.20 % | **26.50 %** | 25.10 dm<sup>3</sup> | 10.78 % | **13.25 %** | 12.88 % |
| abdominales | 21.41 dm<sup>3</sup> | 16.80 % | **17.76 %** | 12.28 % | 18.59 dm<sup>3</sup> | 10.14 % | **11.62 %** | 9.82 % |
| perinealis | 0.53 dm<sup>3</sup> | 16.43 % | 24.48 % | **26.57 %** | 0.36 dm<sup>3</sup> | 10.03 % | **11.18 %** | 10.34 % |
| deltoidea | 3.38 dm<sup>3</sup> | 9.72 % | 15.69 % | **29.89 %** | 1.83 dm<sup>3</sup> | 16.14 % | 23.84 % | **26.92 %** |
| brachialis | 3.57 dm<sup>3</sup> | 10.22 % | **17.78 %** | 17.49 % | 2.80 dm<sup>3</sup> | 11.68 % | **13.73 %** | 8.96 % |
| cubitalis | 1.74 dm<sup>3</sup> | 14.46 % | **20.98 %** | 18.83 % | 1.67 dm<sup>3</sup> | **15.90 %** | 14.85 % | 9.16 % |
| antebrachialis | 2.26 dm<sup>3</sup> | 17.32 % | **22.80 %** | 15.85 % | 2.11 dm<sup>3</sup> | **8.60 %** | 8.47 % | 3.00 % |
| manus | 1.24 dm<sup>3</sup> | **22.70 %** | **23.02 %** | 21.27 % | 0.93 dm<sup>3</sup> | **12.48 %** | 11.63 % | 7.11 % |
| coxae | 12.87 dm<sup>3</sup> | **17.82 %** | 15.99 % | 16.23 % | 9.56 dm<sup>3</sup> | 10.56 % | **11.73 %** | 11.47 % |
| femoris | 18.98 dm<sup>3</sup> | **12.66 %** | 10.47 % | 2.96 % | 13.00 dm<sup>3</sup> | **8.02 %** | 7.82 % | 4.78 % |
| genus | 3.40 dm<sup>3</sup> | 13.76 % | **16.47 %** | 9.59 % | 3.57 dm<sup>3</sup> | 7.94 % | **8.66 %** | 3.46 % |
| cruris | 5.63 dm<sup>3</sup> | 12.17 % | **16.24 %** | 5.86 % | 5.53 dm<sup>3</sup> | **9.02 %** | 8.62 % | 2.88 % |
| pedis | 2.67 dm<sup>3</sup> | **23.06 %** | **22.82 %** | 19.47 % | 1.92 dm<sup>3</sup> | **11.25 %** | 9.51 % | 4.46 % |

**Figure 3.**
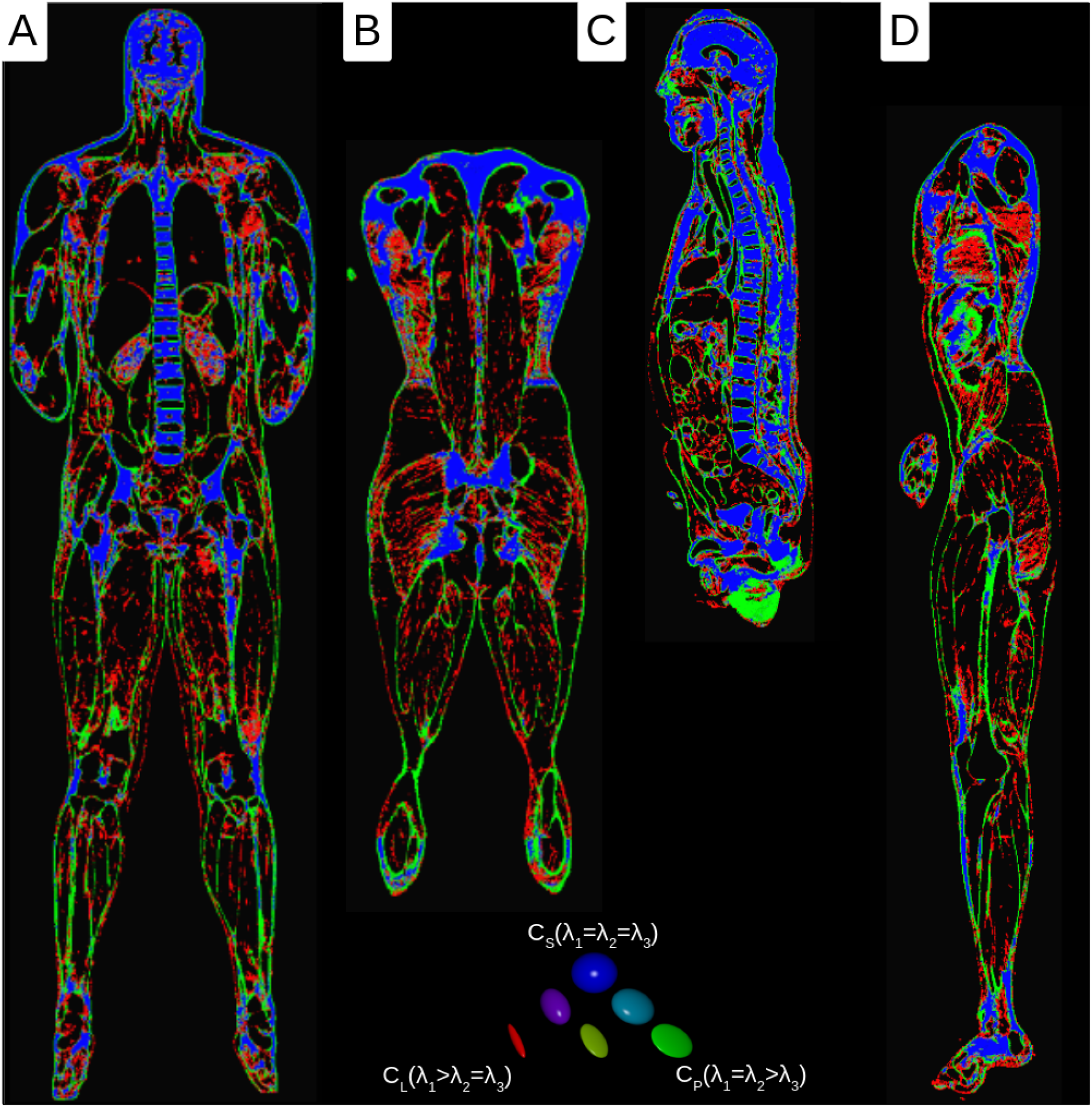
Westin plots of the tensor data of the male body donor (red = fibrous, green = planar, and blue = spherical parts). A) medial frontal section B) dorsal frontal section C) sagittal section D) parasagittal section

**Figure 4.**
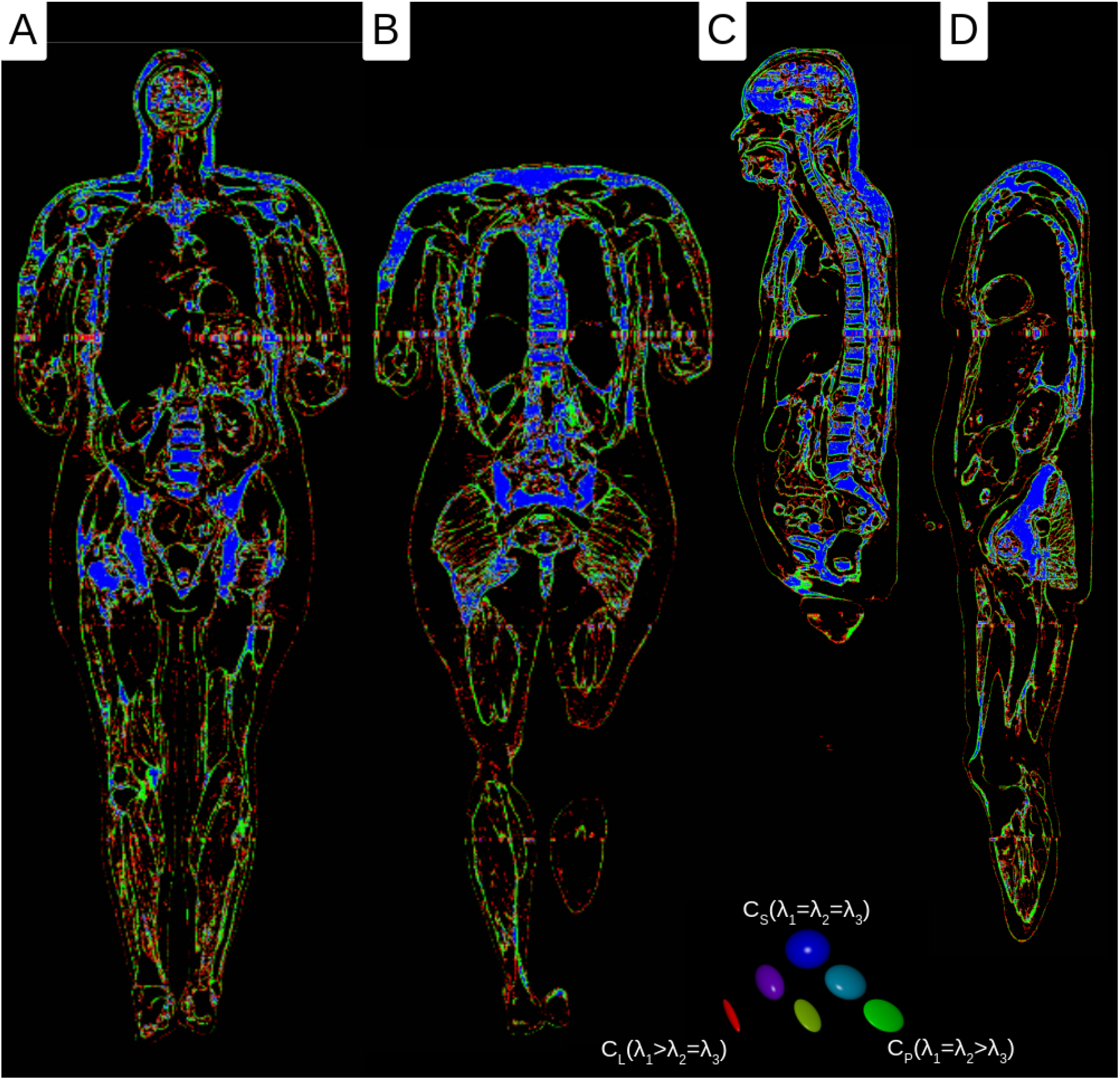
Westin plots of the tensor data of the female body donor (red = fibrous, green = planar, and blue = spherical parts). A) medial frontal section B) dorsal frontal section C) sagittal section D) parasagittal section

### Data Records

#### Connective tissue

The baseline data sets for the male and female body donor, taking into account connective tissue (fig. 2), consist of two data sets with original resolution and reduced bit depth (male & female). Isometric voxel data were created from these data sets by summative binning (male-iso & female-iso). For a quick overview, additional millimetre resolution data sets were calculated from isometric voxel data by summative binning (male-small & female-small).

- male.nii.gz (3753×2141×1867 unit8 – 0.144×0.144×1 mm)
- male-iso.nii.gz (1081×617×3734 uint16 – 0.5×0.5×0.5 mm)
- male-small.nii.gz (540×308×1867 uint16 – 1×1×1 mm)
- female.nii.gz (4096×3061×5190 uint8 – 0.144×0.144×0.33 mm)
- female-iso.nii.gz (1787×1336×5190 uint16 – 0.33×0.33×0.33 mm)
- female-small.nii.gz (590×441×1713 uint16 – 1×1×1 mm)

#### Body regions

To categorise the available data concerning their distribution in the body, an anatomical classification was used^23^. For this purpose, a mask data set was created with 3D Slicer (Brigham and Women’s Hospital (BWH), USA. URL: https://www.slicer.org)^24^ for both men and women, and individual body regions were assigned (figs. 5 and 6). We do not expect a loss of accuracy from using the reduced dataset because the anatomical categorisation was carried out at a low level of detail.

**Figure 5.**
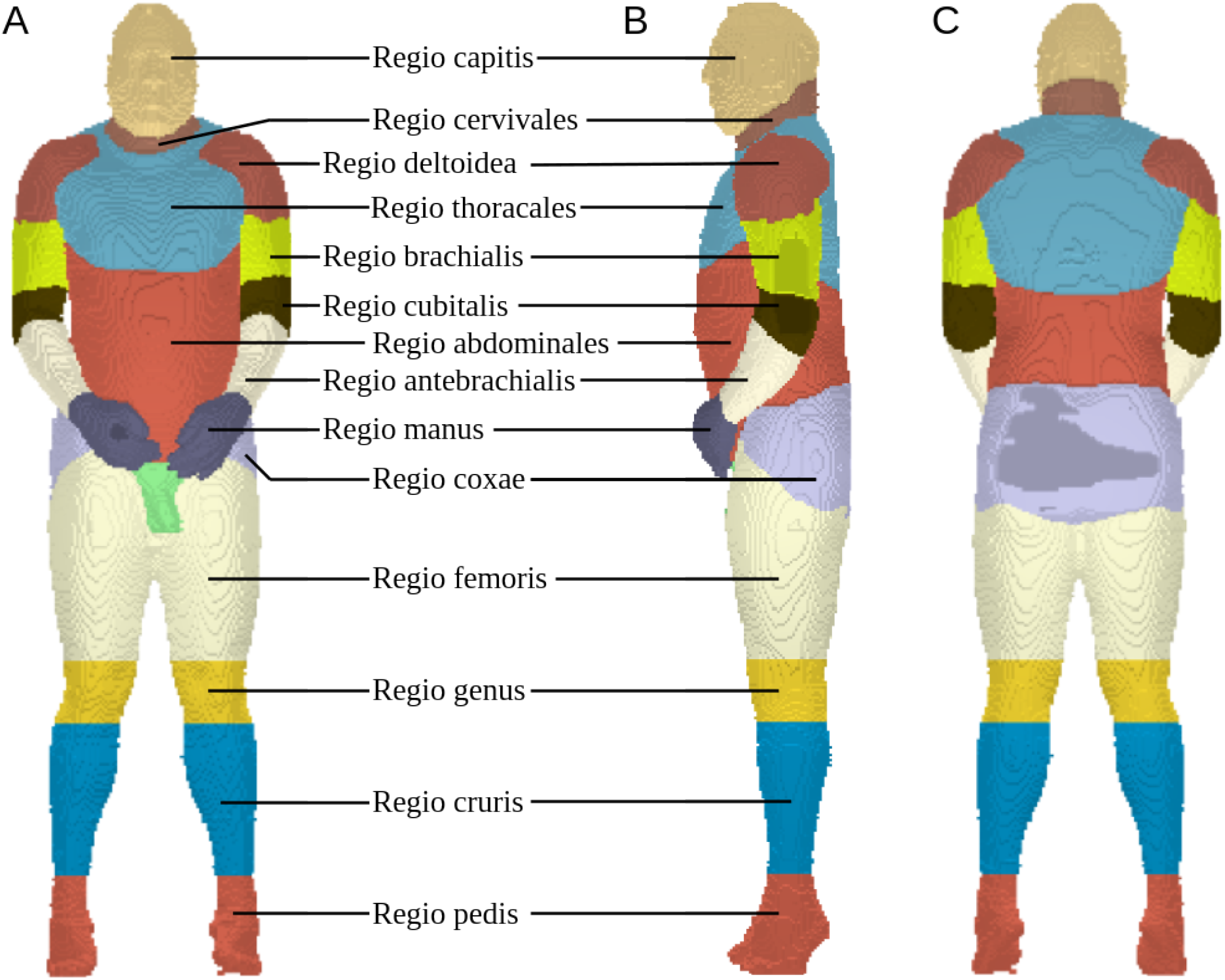
Segmentations of the body regions for the male body donor in random colours. A) Frontal view B) Lateral view C) Dorsal view

**Figure 6.**
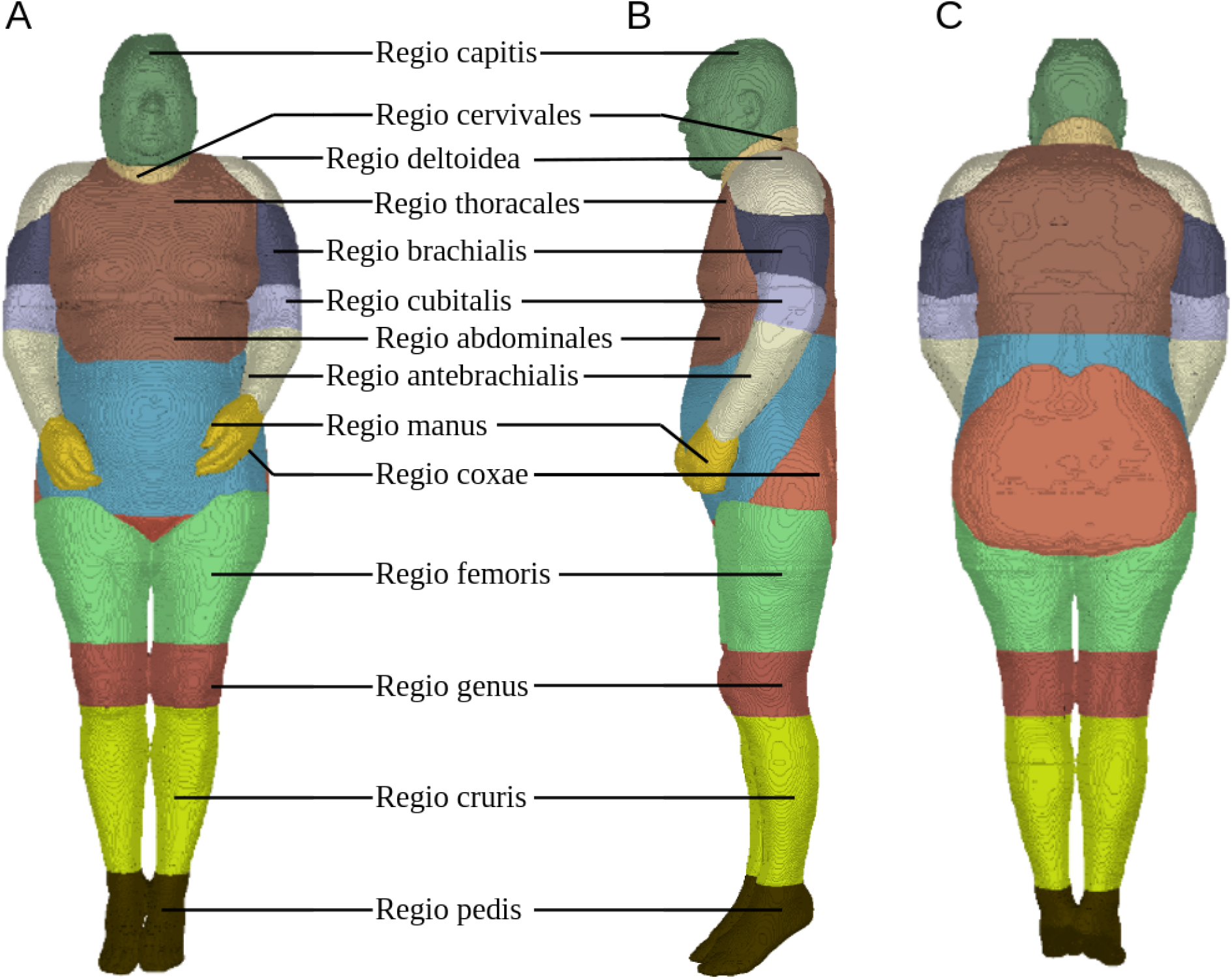
Segmentations of the body regions for the female body donor in random colours. A) Frontal view B) Lateral view C) Dorsal view

- male-small.labels.nii.gz (108×62×373 uint16 – 5×5×5 mm)
- female-small.labels.nii.gz (295×220×856 uint16 – 2×2×2 mm)

#### Thickness measurement

The thickness measurement showed a very variable distribution of connective tissue in terms of its areal thickness in the body (fig. 2). Values between 0.5 and 32 mm were measured for the entire data set. Values above 32 mm can also occur, but these were neglected here due to the time required. The thickest layers of connective tissue are found in the knees, shoulders, and head (table 1). The musculature usually shows very thin or fibrous structures (table 1 *Regio femoris/cruris*). It should be emphasised that the data show a shirt-like distribution of the superficial connective tissue in the thoracic region.

- male-iso.t.nii.gz (1081×617×3734 uint – 0.5×0.5×0.5 mm)
- female-iso.t.nii.gz (1787×1336×5190 uint – 0.33×0.33×0.33 mm)

#### Orientation measurement

The result of the local orientation tensors showed that the highest percentage of local orientation tensors within the fibrous parts was found in the thigh (table 2). The planar portions predominate, except for the head, where the spherical portions predominate. It should be noted that the scan radius for determining the anisotropy was only 5 mm. Thus, larger structures (for example, the brain) are generally classified as spherical, although they could also be planar when viewed on a larger scale.

- male-iso.c2.nii.gz (540×308×1867 symmat – 1×1×1 mm)
- male-iso.c2_e1.nii.gz (540×308×1867 rgb – 1×1×1 mm)
- male-iso.c2_e2.nii.gz (540×308×1867 rgb – 1×1×1 mm)
- male-iso.c2_westin.nii.gz (540×308×1867 rgb – 1×1×1 mm)
- male-iso.c2_westin_rank.nii.gz (540×308×1867 rgb – 1×1×1 mm)
- female-iso.c2.nii.gz (893×668×2595 symmat – 0.66×0.66×0.66 mm)
- female-iso.c2_e1.nii.gz (893×668×2595 rgb – 0.66×0.66×0.66 mm)
- female-iso.c2_e2.nii.gz (893×668×2595 rgb – 0.66×0.66×0.66 mm)
- female-iso.c2_westin.nii.gz (893×668×2595 rgb – 0.66×0.66×0.66 mm)
- female-iso.c2_westin_rank.nii.gz (893×668×2595 rgb – 0.66×0.66×0.66 mm)

### Technical Validation

The data for segmented connective tissues are compared to the scientific record to assess the accuracy of the data. First, we compare the Achilles tendon.

The Achilles tendon is largely visualised in the female dataset as a compact strand with little internal structure. Occasional lines of voxels not marked as connective tissue are a vague indicator of fascicle orientation. Only in the proximal tendon close to the myotendinuous junction substrands can it be identified. They fan out towards the proximal end of the tendon. The diameters of these substrands can be narrowed down to 150 µm to 450 µm so that they most probably correspond to fascicles, which have diameters of 50 µm to 400 µm in human Achilles tendon^25^. We conclude that individual fascicles and subtendons can be resolved when isolated but not within compact regions of connective tissue. The single fibres (diameters of 10 µm to 50 µm) are below the resolution of the dataset. Tensor analysis renders the Achilles tendon as a connective tissue with a three-dimensional extent. However, tendons and ligaments should be identified as unidirectional tissues^26^. Consequently, the diameter of the tendon seems to exceed the threshold for identification as a tissue of unidirectional extent. However, the fascicle orientation is not resolved with sufficient clarity to contribute to the dimensionality result. In the distal part of the Achilles tendon, the Plantaris tendon runs superficially parallel to it and is pressed against the Achilles tendon along parts of their common course^27^. The distinction of both tendons in this region is difficult. We could not judge whether both tendons run parallel or merge from our data set.

Collagen fibres are known to be continuous from the tendon proper through the unmineralised fibrocartilage up to the depth of the mineralised fibrocartilage in the Achilles tendon insertion^28,29^. However, our data set only shows connective tissue up to the mineralisation front. Collagen within the bone and mineralised fibrocartilage appear masked to the methods used. At the posterior end of the Calcaneus, where the Achilles tendon is pressed against it, periosteal fibrocartilage is known to occur^30^. It depicts the methods used here. In conclusion, connective tissues that contain more type II collagen are also rendered^29^.

### Usage Notes

All data are in a compressed NIFTY format (Neuroimaging Informatics Technology Initiative). Common volume data processing software (Amira, 3D Slicer, or Imagexd) can read and process the data. Limitations exist only concerning the available working memory (RAM). They are available for download under Mendeley Data:

- Male DOI: 10.17632/zc53h3dcfg.1
- Female DOI: 10.17632/7m9z78jpgs.1

## Code availability

All software used here is freely available. In particular, the scripts used here for ‘Imagexd’ are freely available open source in the GitHub repository: https://github.com/heikostark/VHP-classification

## Acknowledgements

We thank the U.S. National Library of Medicine for providing the data from the Visible Human Project®. We would also like to thank Stefan Schuster, Irina Mischewski, and the KIP Fascia Research Group (Especially: Christian Puta, Martin S. Fischer, and Robert Schleip) for stimulating discussions.

## Author contributions statement

H.S. segmented geometries, prepared data files, and completed the final processing steps for the 3D data. H.S. and J.S. analysed the results and contributed to writing the manuscript. All authors reviewed the manuscript.

